# Demographic Inferences and Loci Under Selection in a Recently Expanded Coral Population

**DOI:** 10.1101/2021.06.17.448885

**Authors:** James E. Fifer, Nina Yasuda, Takehisa Yamakita, Sarah W. Davies

## Abstract

Coral poleward range expansions in response to warming oceans have been historically observed, however contemporary expansion rates of some coral species have become more rapid as global temperatures rise at unprecedented rates. Range expansion can lead to reduced genetic diversity and surfing of deleterious mutations in expanding populations, potentially limiting the ability for adaption and persistence in novel environments. Successful expansions that overcome these founder effects and colonize new habitat have been attributed to multiple introductions from different sources, hybridization with native populations, or rapid adaptive evolution. Here, we investigate population genomic patterns of the reef-building coral *Acropora hyacinthus* along a latitudinal cline that includes a well-established range expansion front in Japan using 2b-RAD sequencing. A total of 184 coral samples were collected across seven sites spanning from ∼24°N to near its northern range front at ∼33°N. We uncover the presence of three cryptic lineages of *A. hyacinthus*, which occupy discrete areas within this region. Only one lineage is present at the expansion front and we find evidence of its historical occupation of marginal habitats. Within this lineage we also find evidence of bottleneck pressures associated with expansion events including higher clonality, increased linkage disequilibrium, and lower genetic diversity in range edge populations compared to core populations. Asymmetric migration between populations was also detected with lower migration from edge sites. Lastly, we describe genomic signatures of local adaptation potentially attributed to lower winter temperatures experienced at the more recently expanded northern populations. Together these data illuminate the genomic consequences of range expansion in a coral and highlight how adaptation to colder temperatures along the expansion front may facilitate further range expansion in this coral lineage.

## Introduction

Populations are often locally adapted to specific environmental conditions that allow them to thrive. Large-scale environment changes influence these niche conditions and can result in expansion, contraction, or extinction of populations or entire species [1]. Poleward expansion in response to warming oceans is a recent trend, with some species increasingly found in higher latitudes (also referred to as tropicalization), specifically in regions where habitats are degraded or altered [2]. These expanded populations can create socio-political tensions between states and even across countries as commercially relevant species – and species that provide habitats for them – modify their ranges without respect for political and sociological boundaries [3].

An expanding population can experience strong neutral and non-neutral processes. Neutral founder effects can randomly change allele frequencies [4], [5] and low frequency alleles can thereby ‘surf’ on the wave of expansion leading to high frequencies at the range front [6], [7]. These neutral processes can lead to reductions in genetic diversity [8]–[10], higher mutational load if deleterious alleles surf [11], and false signatures of selection [5], [6], [12]. In contrast, non-neutral effects associated with range expansion can produce strong selective pressures through spatial sorting [13], where colonizers of new edge populations have high dispersal abilities [10], and/or natural selection when there is some fitness benefit to being an early colonizer [15]. Additionally, expansion into new environments implies that populations are exposed to novel selection pressures; for poleward expansion, this entails selection pressures imposed by different temperature regimes (i.e. increased seasonality) [16], [17].

Understanding how an organism’s range responds to environmental change is particularly important for species that are biological pillars for essential ecosystems, as corals are to coral reefs. Corals are some of the world’s most important habitat-forming marine organisms, however their recent declines are global in scale [18]. Their sensitivity to temperature [18] and rapid dispersal capabilities (routinely >10km/year; reviewed in Jones et al [19]) mark corals as candidates for range expansion under predicted future warming [20]. The survival of coral species under a changing climate might depend on their ability to successfully shift their ranges; however, studies examining coral range expansion remain scarce [21], [22]. In Japan, the coral *Acropora hyacinthus* has been observed to expand its range [23], which has been correlated with rising temperatures in the region, with temperature increases of 0.5°C/decade between 1982 and 2010 [24]. These increases are likely mediated by the nearby Kuroshio Current’s high sea surface temperature (SST) warming rate (two to three times faster than the global mean SST warming rate) [25]. This ocean warming has promoted macroalgal-to-coral phase shifts both directly, by increased competition from the expansion of tropical corals into the contracting temperate macroalgae range, and indirectly, via deforestation by the expansion of tropical herbivorous fish [26]. However, historically (∼5,000 years ago) corals existed even further north than their current range limits when temperatures were 2-4°C higher than today [27], suggesting that, like other organisms in this region [28], corals may contract and expand along coastal Japan in response to glacial cycles.

Recent studies focusing on *A. hyacinthus* in Japan have provoked intriguing questions related to their population genetic patterns. Using nuclear and mitochondrial markers, Suzuki et al. [29] found that recently expanded populations showed reduced genetic diversity, which is expected along an expansion front [30]. Additionally, they discovered the presence of four genetically distinct *A. hyacinthus* lineages in the region. Only one of these lineages (HyaD) existed within the temperate region, while all four were found in the southern Ryukyus with HyaD becoming progressively rarer in southern islands. Nakabayashi et al. [31] found three genetically distinct lineages of *A. hyacinthus* using microsatellite markers and confirmed the presence of only one *A. hyacinthus* lineage within the temperate region. This divergence between mainland Japan and the Ryukyus has also been observed for a number of terrestrial (reviewed in [32]) and marine species [33]–[42] between the terrestrial biogeographical barrier called the Watase line (also referred to as the Tokara gap). This temperate-subtropical genetic separation has been attributed to allopatric speciation during glacial maximums [39], [41]–[43], the opening and closing of oceanographical barriers facilitated by shifts in the Kuroshio current during glacial maximums [34], [35], [40], [43], and divergent environmental selection pressures [36], [37], [44].

Here, we examine populations of the coral *A. hyacinthus* near their range edge surrounding mainland Japan, which includes a recently colonized site (∼40 years; [23]), and nearby populations in the subtropical Ryukyus islands (Fig. 1). Using population genomics, we 1) identify *A. hyacinthus* cryptic lineages and their associated distributions and demography, 2) further explore the recent range expansion and core-marginal population genomic dynamics within the temperate region, and 3) identify potential loci under selection on the range expansion front that might confer local adaptation to differential thermal regimes experienced across the range.

**Figure 1.**
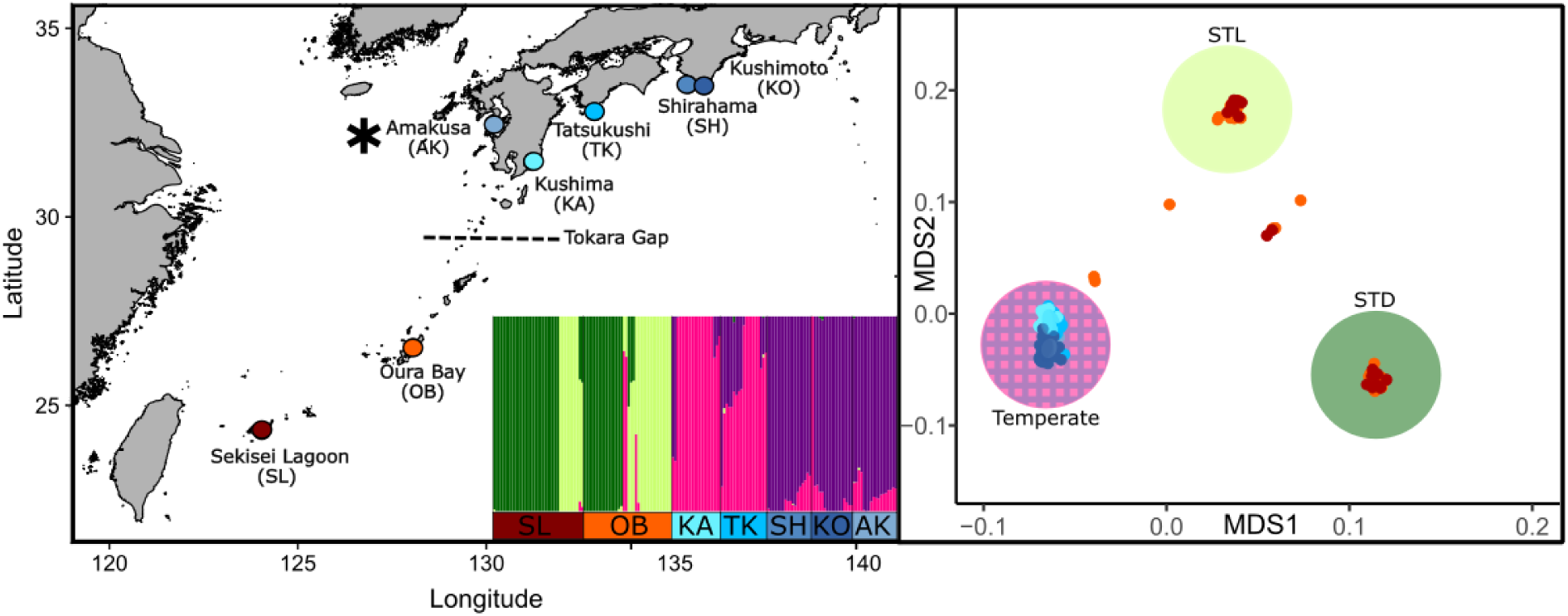
Left panel: Map of all coral collection sites in Japan including site names (abbreviation). Asterisk denotes recently expanded site (in the last 40 years), Amakusa. Inset: Results from Admixture analysis where each bar represents an individual coral colony and each color denotes its inferred membership to each of the 4 putative ancestral populations. Right panel: Multidimensional scaling (MDS) plot based on genetic covariance matrices of all samples, which highlights the presence of three major cryptic lineages (STL, STD, and Temperate).

## Methods

### I. Sample collection, sequencing and pre-processing

#### Coral colony sampling and 2b-RAD-seq library preparations

Fragments of *A. hyacinthus* colonies (33-50 colonies/site between 5-300 cm in diameter) were collected from seven sites in Japan between 2009 and 2016 via SCUBA (Supp. Table 1; Fig. 1) under prefectural permit 21-18, 24-36, 25-44 and with agreement from local fishermen’s unions. Samples were immediately preserved in 99% ethanol, stored at −20℃, and transferred to the United States under CITES permit (13JP002699/TE, 14JP000665/TE, T-WA-12-001588, T-WA-13-002421, T-WA-14-00589). DNA was isolated using a modified phenol-chloroform extraction method [45], cleaned with ZYMO-DNA-clean-and-concentrator kits, and resulting extracts were prepared for 2b-RAD-sequencing following Wang et al [46]. Eight replicate samples were prepared to assist with clone identification. A total of 188 samples (19-40 samples per site; Table 1) were successfully barcoded and sequenced across four lanes of Illumina HiSeq 2500, High Output v4 using single-end 50bp at Tufts University Core Facility (TUCF).

#### Genotype calling

Raw reads were trimmed, deduplicated and quality filtered with FASTX TOOLKIT (http://hannonlab. cshl.edu/fastx_toolkit) and only reads with Phred scores > 20 maintained. Quality-filtered reads were mapped to the *Acropora millepora* genome [47] via bowtie2 [48]. Genotyping was performed using ANGSD v0.921 [49]. Standard filtering that was used across all analyses included loci present in at least 80% of individuals, minimum mapping quality score of 20, minimum quality score of 25 (unless no minimum allele frequency (MAF) filter was used in which case quality scores of 25 and 30 were used), strand bias p-value > 0.05, heterozygosity bias > 0.05, removing all triallelic sites, removing reads having multiple best hits and lumped paralogs filter. All analysis pipelines are open source and can be found at https://github.com/jamesfifer/JapanRE.

#### Clone identification

Additional filtering was performed to detect clonemates, which included only sites with MAF > 0.05, depth of coverage > 5 reads, and SNP p-value > 0.05. Clones were detected using hierarchical clustering of samples based on pairwise identical by state (IBS) distances calculated in ANGSD. Technical replicates were used to identify appropriate height cutoff (Supp. Fig. 1). Only one clonemate was retained for all downstream analyses (Supp. Table 1).

### II. *Acropora hyacinthus* Lineage Analyses

#### Lineage assignments

Additional site filtering was performed prior to downstream analyses, which included only sites with MAF > 0.05 and linked loci were filtered out using ngsLD [50] and the provided prune_graph.pl script (max distance 5kb, minimum weight 0.2) to ensure population structure was not driven by linkage. To detect admixture in corals from all seven sites, the program ADMIXTURE (v. 1.3.0; [51]) was used to find the optimal number of clusters (K) with the least cross validation error. Single Nucleotide Polymorphisms (SNPs) were hard called using genotype likelihoods estimated by SAMtools with a SNP p-value < 0.05 for ADMIXTURE input only. Principal Component Analysis (PCA) and admixture barplots were then visualized using genotype likelihood estimates (no SNP p-values). PCAs were performed using a covariance matrix based on single-read resampling calculated in ANGSD and ngsADMIX [52] and admixture results were visualized using the K with the least cross validation error reported from ADMIXTURE.

Individual lineage assignments were based on ancestral population assignments from ngsADMIX results, where ancestral populations were only considered lineages if separation was shown on the PCA first axis. If a sample had an assignment proportion >75% to a single cluster (i.e. bar color), it was assigned to that lineage. All other samples were removed from downstream lineage specific analyses (Supp. Table 1).

#### Analyses of genetic divergence, demographics, introgression, and phylogenetics between lineages

In order to determine genetic differentiation between lineages, ANGSD was used to calculate the site allele frequency (SAF) for each lineage and then realSFS calculated the site frequency spectrum (SFS) for all possible pairwise comparisons. These SFSs were used as priors with the SAF to calculate global F_ST_. Here, only weighted global F_ST_ values between lineages are reported. Heterozygosity was then calculated using genotype likelihoods generated by ANGSD and an expectation maximization algorithm (EM) used by realSFS in a custom script (git@github.com:sbarfield/Yap_Ahyacinthus-.git/heterozygosity_beagle.r). Differences in heterozygosity between lineages was calculated via Welch two sample t-test.

Linked sites were removed from downstream analyses using ngsLD and the provided prune_graph.pl script (max distance 5kb, minimum weight 0.2). ANGSD was used to obtain 100 series of 5 block-bootstrapped SFS replicates, which were averaged to create 100 bootstrapped SFS for each lineage. SFS was polarized using the *A. millepora* genome as an ancestral reference. Multimodel inference in moments [53] was used to fit two-population models (https://github.com/z0on/AFS-analysis-with-moments) and all unfolded models were run on 10 bootstrapped SFS and replicated six times. The best fit model was then selected based on lowest AIC value. Parameters (i.e. migration, epoch times, and effective population sizes (Ne)) for the best fit model were obtained by running the best fit model on 100 bootstrapped SFS and replicated six times. Additionally, we ran the unsupervised analysis StairwayPlot v2 [54] to one dimensional SFS as a second effort to reconstruct effective population sizes. For all demographic analyses we used a mutation rate of 4e^−9^ per base per year and generation time of 5 years from estimates for *A. millepora* [55], [56].

Phylogenetic analyses of the 2b-RAD-seq data set were conducted using RAxML [57]. First we explored a range of thresholds for loci coverage between samples (50, 60, 70, 80 and 90% of individuals), MAF filter (none, <0.001, <0.01, <0.05), and minimum depth of coverage (1x, 2x, 4x, 6x, 8x, 10x). RAxML, with a GTRGAMMA model and 100 rapid bootstraps, was used to explore the resulting concatenated sequence matrices and choose filters that produced the most reliable trees (based on node resolution and support). All trees agreed in terms of separation of the three lineages and we used the standard filtering from other analyses (80% of individuals, minimum 1x depth of coverage) plus a MAF filter < 0.05 because this maximized bootstrap support.

#### Identifying loci under selection across lineages

Additional filtering of loci was conducted prior to outlier analyses, which included SNP p-value e^−5^ and MAF < 0.05. Two outlier detection programs were used. First, PCAdapt (v. 4.3.3 [58]) was used to determine the optimal K for all pairwise comparisons using a score plot displaying population structure. After filtering out two outlier individuals that were skewing clustering, a K of 2 was selected for all pairwise comparisons between all lineage pairs. A q-value of 0.05 was used as a cutoff for determining outlier loci. BayeScan (v. 2.1[59]) was then used as a second approach to identify outlier loci. The same two outlier individuals were removed and the F_ST_ outlier method implemented in BayeScan identified outlier loci for each pairwise lineage comparison using 5,000 iterations, 20 pilot runs with length 5,000, and burn-in length of 50,000. We employed the default prior odds of neutrality (10) and a q-value cut-off of 0.05. If the same locus was identified across both analyses, the locus was considered a true outlier and the annotated genes 1kb upstream or downstream of this outlier locus were reported. These genes were then compared with the module of genes previously associated with the environmental stress response (ESR) in Pacific *Acropora* corals [60].

### III. Analyses Within the Recently Expanded Temperate *Acropora hyacinthus* Lineage

Only samples from the northern sites surrounding mainland Japan (Amakusa, Kushima, Kochi, Kushimoto, and Shirahama; Fig 1), which were assigned to the temperate lineage (N= 93; Supp. Table 1) were used for the following analyses.

#### Population genetic structure, expansion direction, testing theoretical expansion consequences and demographics

To investigate basic population genetic structure within the temperate lineage, we implemented PCAs and admixture analyses, and calculated global weighted F_ST_ for all pairwise comparisons and expected and observed heterozygosities for each population. All analyses were conducted as described within Section II: *Acropora hyacinthus* lineage analyses. The PERMANOVA function *Adonis* (from the R package vegan [61]) was used to determine if there was clustering according to core or edge. To test for differences in the number of clone pairs between range edge and core populations, a Pearson’s chi-squared test was used.

Next, Slatkin’s directionality index Ψ [62] was calculated to confirm the direction of expansion. Loci with fixed ancestral alleles were removed using SAMtools [63] and the directionality index Ψ was calculated using the script devtools::install_github(“BenjaminPeter/rangeExpansion”, ref=“package”) and the function get.all.psi. Linkage disequilibrium (LD) was estimated independently for each population using ngsLD [50], and r^2^ was plotted for each population using fit_LDdecay.R (max 200kb).

Purifying selection was assessed to estimate the degree of mutational load. To calculate purifying selection, sites with synonymous and missense mutations were identified using Variant Effect [64]. Global Watterson’s theta was calculated for each site using the thetaStat tool in ANGSD from the maximum likelihood estimate of the SFS. Watterson’s theta was estimated at this subset of sites for all temperate populations. We then correlated the ratio of missense to synonymous site diversity with the site’s distance from Kushima using Pearson’s correlation coefficient. Lastly, demographic analyses in moments were run as described for the coral lineage analyses between all five temperate populations surrounding mainland Japan.

#### Identifying selection pressures and loci under selection

Patterns of isolation-by-distance (IBD) were evaluated among the temperate populations using a Mantel test with 10,000 random permutations between the F_ST_ matrix and pairwise oceanographic distance. A Mantel test was also performed between the F_ST_ matrix and pairwise differences in monthly minimum, maximum and mean SST. These SST data were extracted from the Modis Satellite (https://oceancolor.gsfc.nasa.gov/).

Outlier analyses were conducted as described previously between the two ancestral populations within the temperate region identified using ADMIXTURE and ngsADMIX.

## Results

### Presence of cryptic *Acropora hyacinthus* lineages with distinct evolutionary histories

Analyses of population structure with ADMIXTURE and ngsAdmix using samples from all sites show the presence of four (K=4) ancestral populations with the majority of individuals assigning with high proportion (greater than 0.75) to a single population (Fig. 1). PCoA based on pairwise genetic distances demonstrates that these populations can be separated into three distinct clusters (referred to as lineages here) (Fig. 1). All three lineages are found in Oura Bay, two are present in Sekisei lagoon and only one is present in the temperate region. Hereafter these are referred to as subtropical light green (STL) subtropical dark green (STD) and temperate lineages.

Pairwise F_ST_ indicated high genetic differentiation (0.1770 - 0.236) between all lineages (Supp. Fig. 2A) and phylogenetic analyses confirmed the presence of three phylogenetically distinct clusters with high bootstrap support (Supp. Fig. 2B). Expected heterozygosity was lower in the temperate lineage compared to the other two lineages (p < 0.05) (Supp. Fig. 2C)

Demographic analyses for all three pairwise comparisons showed best fit models included a scenario supporting the presence of genomic islands (Supp. Fig 3; Supp. Table 2). All analyses between all three pairwise lineage comparisons showed support for a split roughly 100 kya (Fig. 2). Both subtropical lineages (STL, STD) showed asymmetrical migration with the temperate lineage with higher gene flow from subtropical lineages to the temperate lineage. Migration between subtropical lineages was symmetrical and it is also worth noting that STD and the temperature lineage experienced an epoch with no migration (Fig. 2; Supp. Fig. 4). Effective population sizes were estimated to be consistently lower in the temperate lineage when compared to the two subtropical lineages (Fig. 2). Stairway plot analysis was primarily consistent with the moments analysis, with the exception of the appearance of recent Ne contractions in the past few thousand years with Ne increases starting roughly 50 kyr earlier for all three lineages (Fig. 3). It is noteworthy that the temperate lineage was the first lineage to exhibit this contraction.

**Figure 2.**
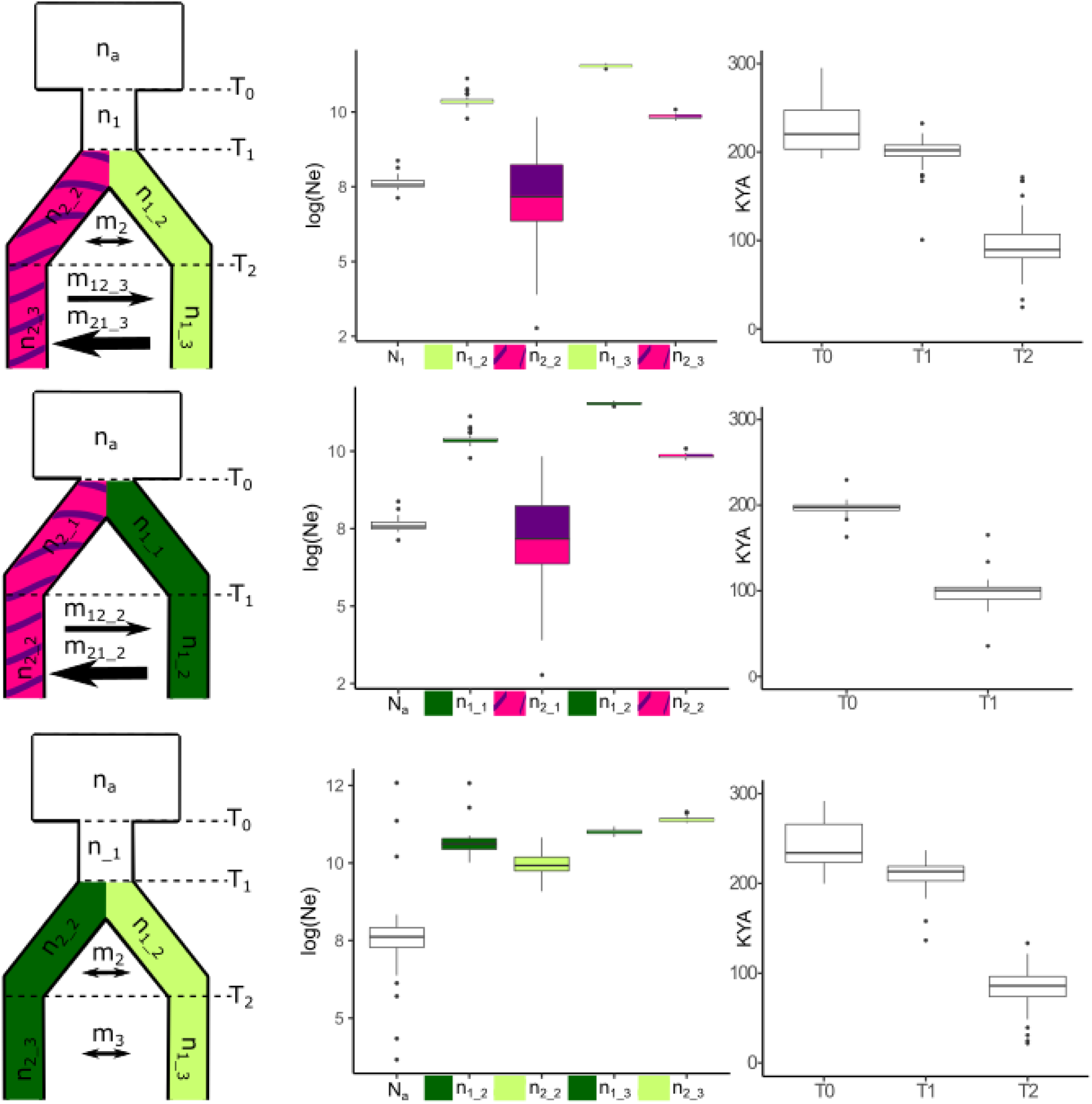
Moments parameter output for pairwise lineage analyses. n=effective population; n1_1=population 1, epoch 1; m=migration. Arrow size corresponds to migration rate in case of asymmetrical migration. Boxplots display mean log(Ne) and mean time since introgression in kya (thousand of years from present) with the box representing the interquartile range (IQR) between the upper and lower quartile. The whiskers extend from the hinge to the highest value that is within 1.5 * IQR of the hinge.

**Figure 3.**
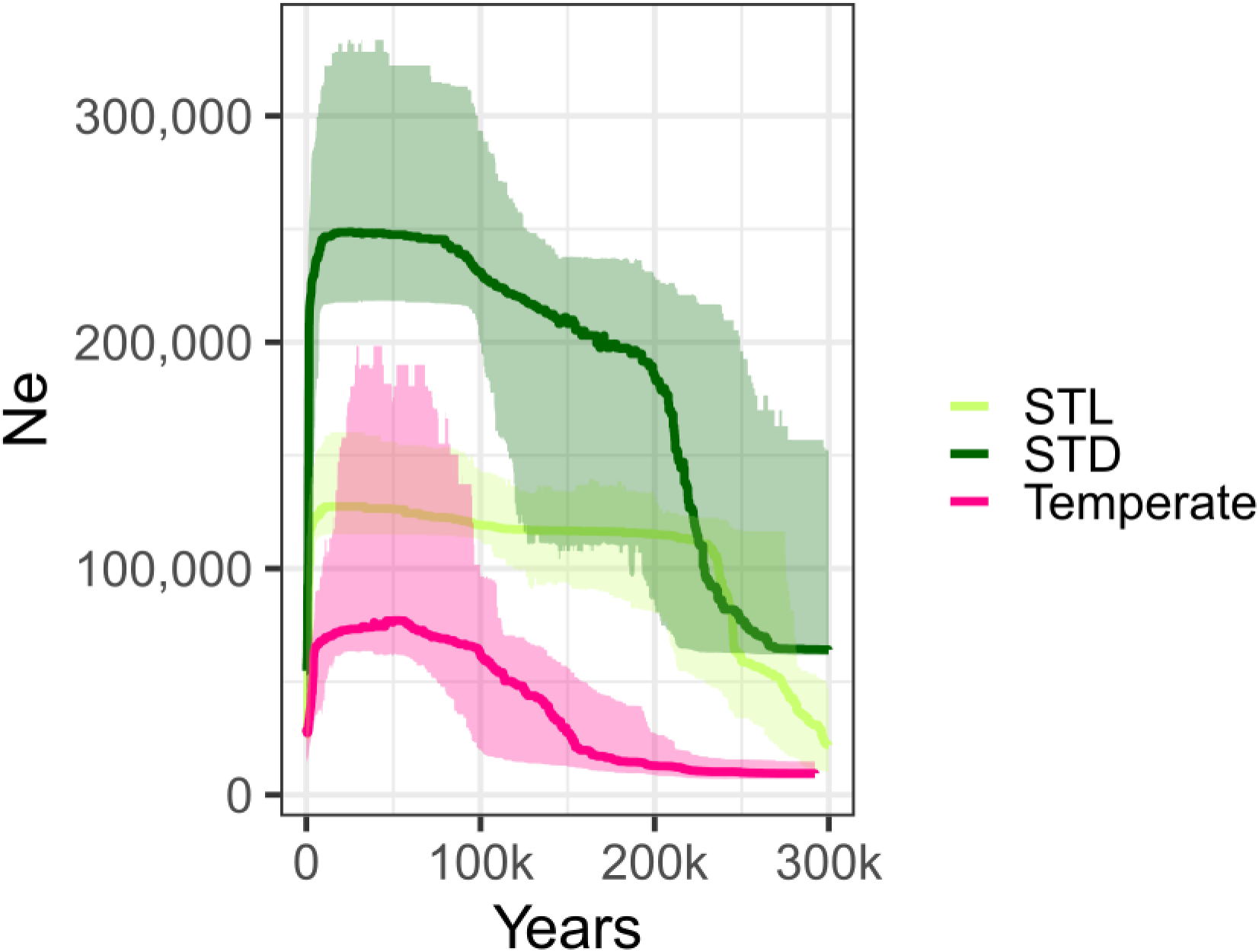
Stairway Plot showing effective population sizes (N_e_) through time for each *Acropora hyacinthus* lineage. Lines show median N_e_, ribbons represent 95% confidence intervals.

In the STL-temperate lineage comparison we identified outlier loci near 41 genes – 20 of which were in the top three environmental stress response (ESR) gene modules from Dixon et al (2020). In the STD-temperate comparison we found outlier loci near 24 genes, 12 of which were ESR candidates. Five outlier genes overlapped in both temperate-subtropical lineage comparisons, three of which were ESR candidates (centrosomal protein 290kDA, elongation of very long chain fatty acids protein 4 (EVOL4) and one unannotated gene). In the STD-STL comparison we found 56 outlier genes, 23 of which were ESR candidates (Supp. File 1).

### Genetic signature of range expansion and demographics within the temperate lineage

Analysis of population structure with ngaAdmix and pcAngsd demonstrated separation of genetic variation (p < 0.001) between the three edge sites compared to the two core sites (Fig. 4A; Fig. 4B). Weighted F_ST_ estimates demonstrate low levels of differentiation among edge populations (0.017-0.024) and among core populations (0.017) and higher levels of differentiation between edge and core (0.036-0.045) (Fig. 4C). The directionality index Ψ for range expansions [62] supported the more southern sites (Kushima/Kochi) as the ancestral origin for Amakusa, Shirahama and Kushimoto (Fig. 5). Expected heterozygosity was higher in core populations compared to edge populations for all pairwise comparison (p < 0.05) (Fig. 6B). Nucleotide diversity at synonymous and missense sites followed the same trend; however, edge sites showed a greater decrease in nucleotide diversity at missense sites, which resulted in a decreasing thetaN/thetaS ratio as populations were further from the proposed origin site Kushima (Fig. 6C). This result suggests that purifying selection is stronger as distance from Kushima increases (Fig. 6C). While LD was similar across all temperate populations, edge populations did show a trend of higher LD (Fig. 6A). Edge populations were also home to significantly more clone pairs when compared to the core populations (Pearson chi-squared=23.969, df = 1, p = 9.789e-07; Fig. 6B).

**Figure 4.**
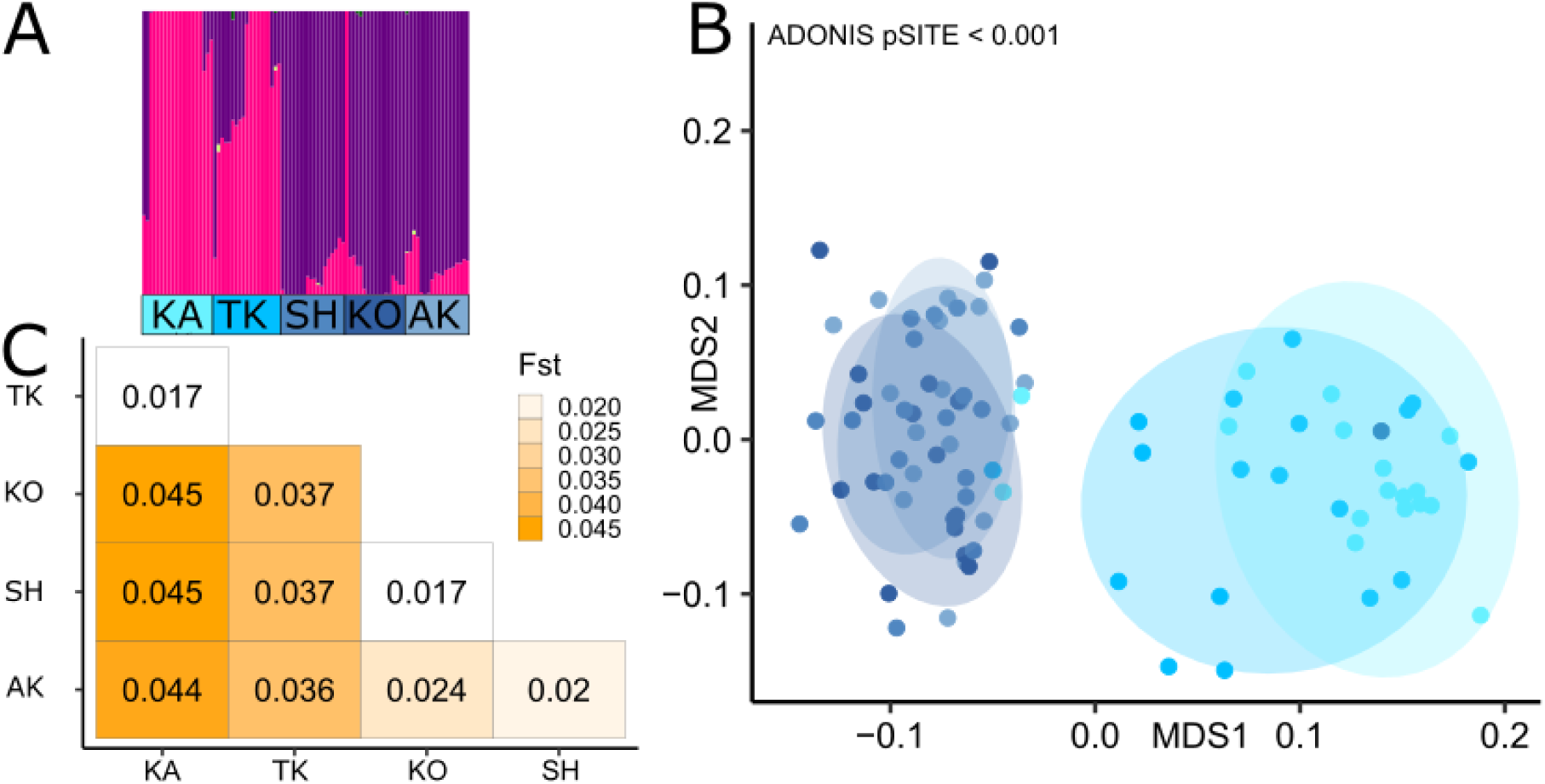
Population genetic structure within the temperate *Acropora hyacinthus* lineage. A) ngsAdmix results where each bar represents a coral individual and the color of the bar represents its inferred membership in each of the 2 potential ancestral populations within the temperate region. B) Multidimensional scaling (MDS) plot based on genetic covariance matrices demonstrating significant clustering between samples from the core sites (light blue) and the edge sites (dark blue). C) Pairwise global F_ST_ between all temperate region sites.

**Figure 5.**
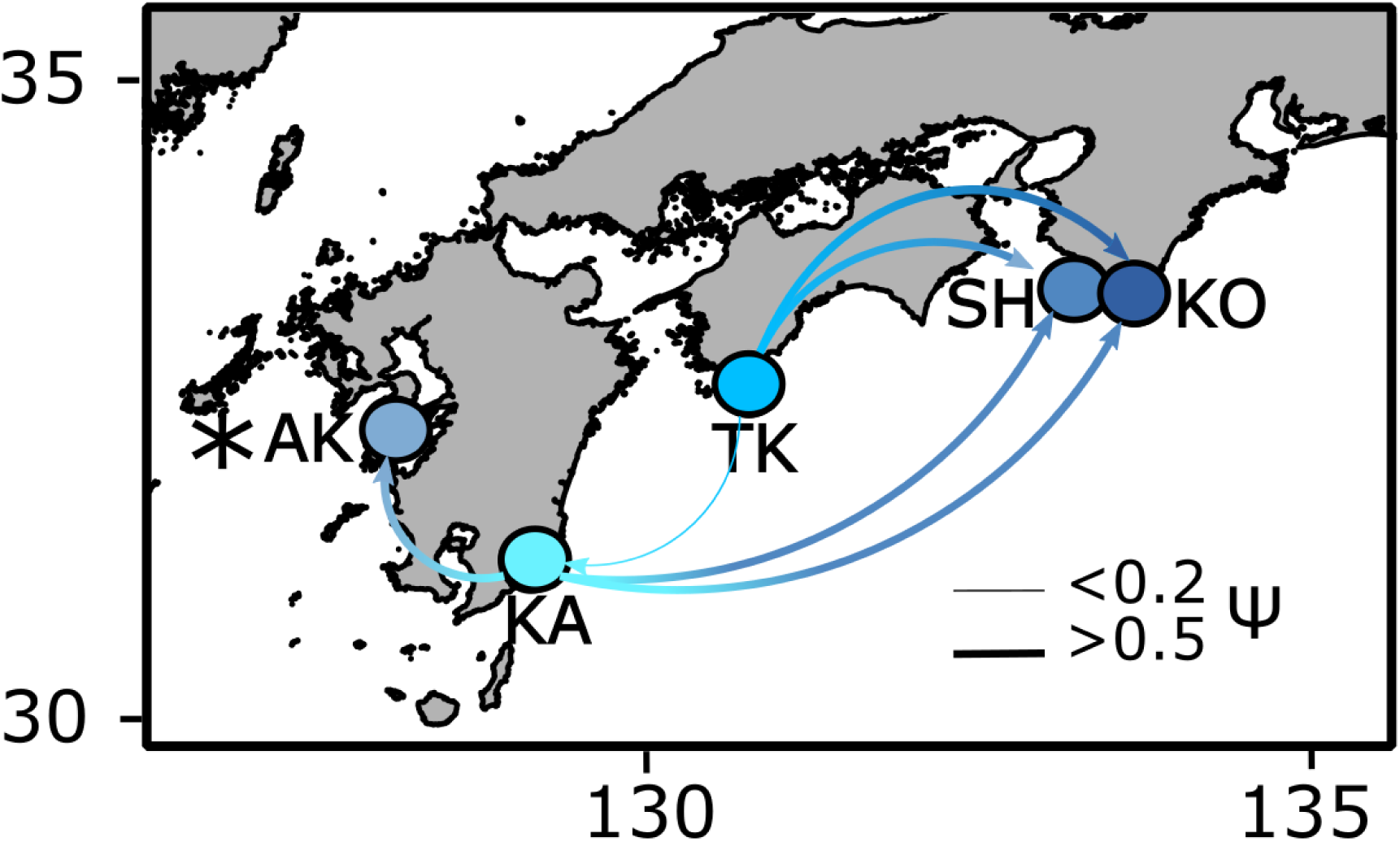
Migration between each temperate lineage *Acropora hyacinthus* population sampled in mainland of Japan, Direction of arrow denotes direction of expansion determined by positive or negative psi value. Thickness of arrow, denoting strength of signal of expansion shows absolute value of psi. Asterix indicates the most recently expanded site that was sampled here, Amakusa (AK).

**Figure 6.**
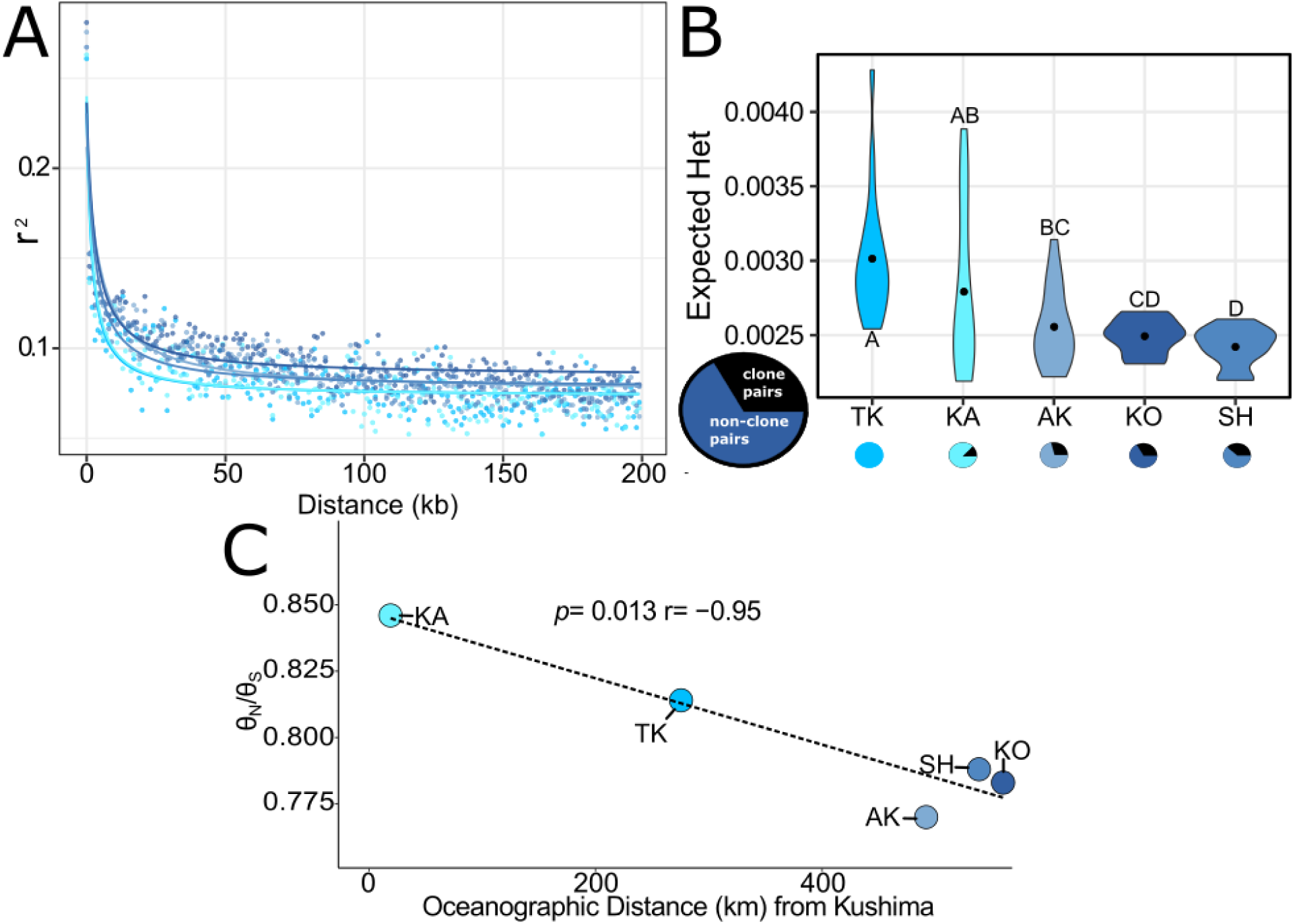
Genetic signatures of range expansion in the temperate *Acropora hyacinthus* lineage. A) Estimated linkage disequilibrium (LD) for 0-200 kb in each populations with line showing best fit decay. B) Mean expected heterozygosity (H_e_) estimates for each population. Letters in common denote non-significant differences (p > 0.05) of H_e_ between sites. Circles below represent the proportion of samples that were made of clone pairs (black) relative to unique genotypes (shades of blue). C) Estimated θ_N_/θ_S_ ratios for each population plotted against oceanographic distance from Kushima (km) with the dotted line representing the linear model fit for purifying selection as distance increases.

Demographic analyses revealed asymmetrical gene flow between all core and edge populations, with lower migration out of edge sites (Fig. 7; Supp. Fig. 5). Migration within edge and within core sites both show symmetrical gene flow (Fig. 7; Supp. Fig. 5) and estimated N_e_ was consistently lower at edge sites compared to core sites (Fig. 7).

**Figure 7.**
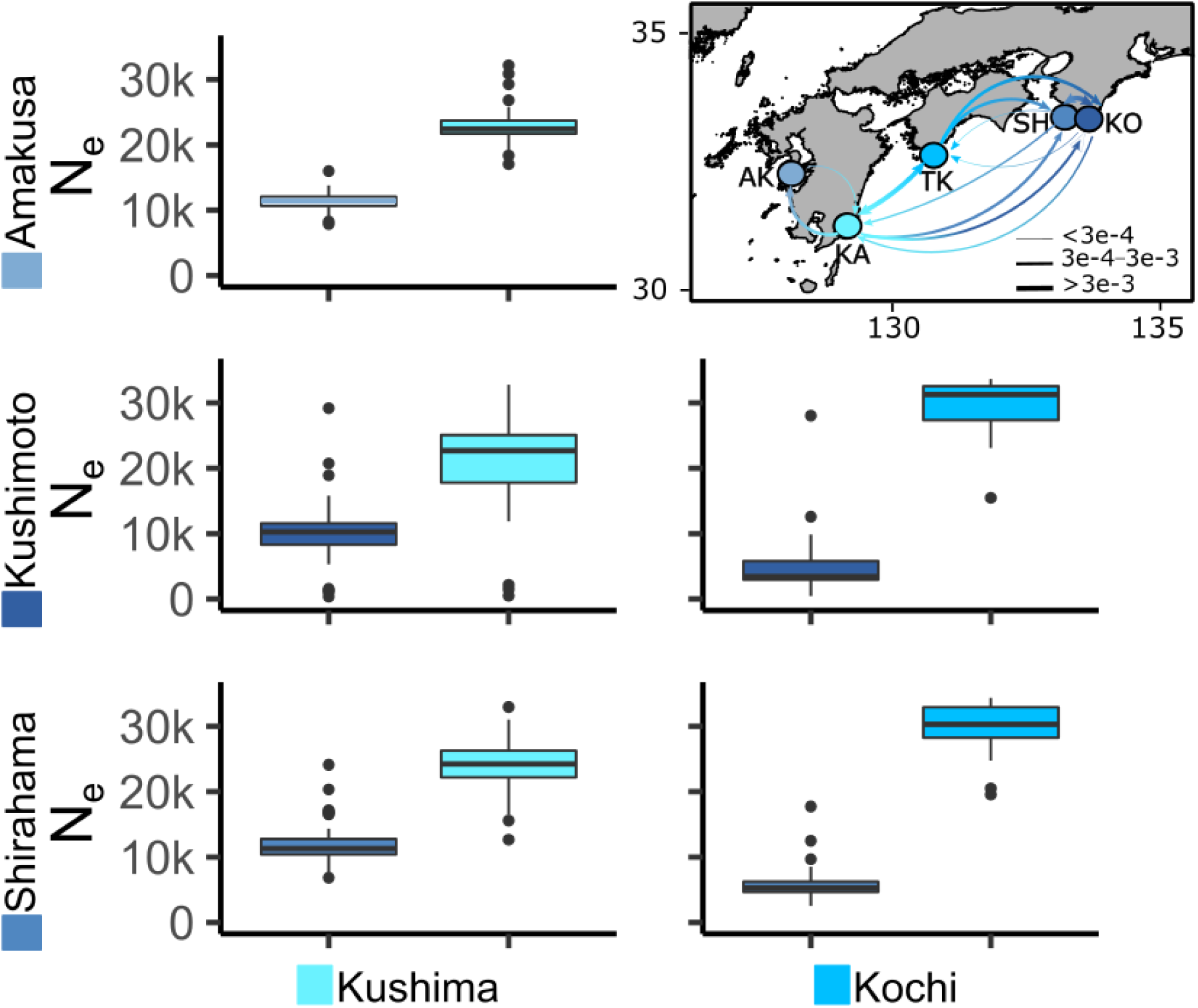
Pairwise moments parameter outputs for the temperate populations of *Acropora hyacinthus* demonstrating mean effective population estimates (N_e_) and mean pairwise migration rates (the fraction of the total population that are new immigrants in each generation) between sites (shown via relative thickness of arrows, double sided arrows represent symmetrical migration).

Mantel test showed non-significant (p > 0.05) signals of isolation by distance (IBD) and non-significant differentiation based on sea surface temperature (SST) minimums, maximums, and means, although mean and minimum SST both showed a higher correlation with F_ST_ (r = 0.2; 0.16) compared to oceanographic distance (r = 0) (Supp. Fig. 6). When comparing edge vs core sites, we detected six genes that were near outlier loci, three of which were ESR candidates (Supp. File 1).

## Discussion

### Presence of *Acropora hyacinthus* lineages that vary in their distributions

Consistent with previous work using different loci (microsatellites [31]; mitochondria and nuclear markers [29]), our 2b-RAD-seq data suggested the presence of three morphologically cryptic *A. hyacinthus* lineages across our sampled sites in Japan. This species is well known for its cryptic lineages with four lineages previously identified in Palau, American Samoa, and Australia, two lineages in Pohnpei and Palmyra [65], two lineages on the island of Yap in Micronesia [66], and two observed in Taiwan [29]. However, the presence of cryptic *A. hyacinthus* lineages is not always observed. For example, Davies et al. [67] found no evidence of cryptic lineages across the entire expanse of Micronesia (even in Palau) using multilocus SSR data. In addition, only one lineage has been reported in the islands of Moorea and Tahiti in French Polynesia [68], which used the same 2b-RAD-seq approach leveraged here. In Japan, latitude appears to be the main driver of the distribution of the temperate lineage, whereas lineages STL and STD are both common in the two southern Ryukyus sites, which is consistent with Nakabayashi et al [31] and Suzuki et al [29].

Although this study did not investigate the physiological differences between *A. hyacinthus* lineages, coral cryptic lineages are known to exhibit differences in thermal tolerance [69] as well as differences in spawn times [70]–[72]. Interestingly, the temperate and one of the subtropical lineages have previously been shown to spawn at the same time (June) in the Ryukyus [29]. Crosses between subtropical and temperate lineages exhibited low fertilization rates but they were semi-compatible [29], suggesting that these lineages maintain some capacity for hybridization – consistent with our admixture and PCA results. Taken together, our findings of cryptic lineages reiterate the importance of first exploring the potential for divergent lineages before investigating population connectivity, calculating speciation rates, and predicting species distributions.

### Demographic history of *Acropora hyacinthus* lineages

Our demographic analyses suggest that divergence times between all three *A. hyacinthus* lineages occurred during the interglacial interval around 200 kya and this divergence was facilitated by their population expansion. This result is contrary to expectation given that divergence of many other species in the region is attributed to isolation in glacial refugia of which there are several in the region: Okinawa Trough, East China Sea and South China Sea [73]. During interglacial periods, Acroporid corals have historically expanded populations due to rising sea levels [74] and have shifted poleward as temperatures rise [75]. It is therefore possible that this expansion into different regions mediated divergence between lineages; however, our divergence times should be interpreted with caution as these numbers are based on an estimated mutation rate. The temperate and STL lineages showed asymmetrical migration following the split suggesting parapatric or sympatric speciation while the temperate lineage and STD showed a period (200-100 kya) of no migration consistent with allopatric speciation. These inferences are corroborated by present day distributions where the temperature lineage and only one of the subtropical lineages coexist in Taiwan [29].

Given the current distributions, estimated divergence time, and N_e_ increases by both subtropical lineages following divergence, we hypothesized that the divergence of the three lineages would have been facilitated by expansion into the Ryukyus. However, fossil records do not support this hypothesis and instead suggest that the increase in N_e_ was due to population growth and divergence conditioned by the introduction of a geographic barrier between the Ryukyus and Taiwan/Philippines. Acroporids were present in Taiwan in the mid-Pleistocene [76] and in mainland Japan in the Holocene (∼5-6 kya) [27], but no information is available specifically showing the presence of *A. hyacinthus* in either of these regions during the Pleistocene. In contrast, there are fossil records for the lower and upper Ryukyus that demonstrate the presence of *A. hyacinthus* as early as mid-Pleistocene (∼400 kya) [77], suggesting that the increase in N_e_ for both subtropical lineages following the split (∼200 kya) shown here does not correspond to an expansion into the Ryukyus. Given fossil records point to fluctuating sea levels driving community composition shifts in the Ryukyus region [77], we propose the explosion in N_e_ 200 kya for the two subtropical lineages likely captures *A. hyacinthus’s* population growth within these reefs, not their introduction. The absence of gene flow between the temperate lineage and STL lineage, lack of increasing N_e_ for the temperate lineage following the split, and the scarcity of the temperate lineage in modern day lower and middle Ryukyus still point to some degree of geographical separation between the temperate and subtropical lineages 200 kya. However, our results cannot elucidate where this initial separation occurred.

An increase in effective population size during the last interglacial period (100 kya) was observed for all three lineages, which aligns with the pattern of major reef development in the Ryukyus that responded to 100 kyr cycles starting in the mid-Pleistocene [78]. The larger increase in N_e_ 100 kya compared to 200 kya corresponds to relatively higher sea levels observed 100 kya [78] and is consistent with data from *Acropora tenuis* in Japan showing increasing N_e_ during the last interglacial period [79]. This result is particularly interesting as *A. tenuis*’s range overlaps with *A. hyacinthus,* while other Acroporids that are not found as far north as these two species show a decrease in effective population sizes during this time period [79]. Reduced frequencies of Acroporids in the fossil record have been observed during the last interglacial period, potentially suggesting that this group of corals experienced large die-offs during this time, especially in the tropics of the Northern Hemisphere due to increasing temperatures [75]. Given that several Acroporids sympatrically distributed with *A. hyacinthus* exhibited decreasing N_e_ during this time period [80], it is possible that these dieoffs led to niche availability, which *A. hyacinthus* subsequently filled. Additionally, according to limited Pleistocene coral fossil records in the Ryukyus, *A. palifera* (which like *A. hyacinthu*s inhabits the shallow upper reef slope) had higher abundances during the mid-Pleistocene than modern day [77]. These patterns are consistent with patterns observed in Caribbean corals where increases in N_e_ during the last interglacial period have also aligned with simultaneous loss of spatially co-occurring species [81].

In the case of the temperate linage, our demographic analyses suggest these populations have historically inhabited marginal areas in the western North Pacific, including reefs around mainland Japan. Following the increase in N_e_ 100 kya, contact was reestablished between the temperate and the STD lineage, which suggests that range expansion was occurring during this time period. Asymmetrical migration coupled with lower gene flow from the temperate lineage to the subtropical lineage, is consistent with a scenario where the temperate lineage is dominating mainland Japan habitats and due to prevailing northward currents, migrants exhibit northward bias. It is therefore possible that the temperate lineage expanded its range into the upper Ryukyus and around mainland Japan during this time. Lower expected heterozygosities in the temperate lineage also suggest historical occupation of these marginal areas. Limited coral fossil records surrounding mainland Japan can only corroborate presence of reefs as far back as the Holocene [27]. However, cold temperate kelp species – the major competitors for substrate space around mainland Japan – have shifted their range under interglacial/glacial cycles [82], suggesting the mechanism of modern range expansion in this region could have facilitated historical range expansion of *A. hyacinthus* as well. Contractions shown in the Stairway Plot analyses during the last thousand years could be due to global cooling, which started 8-10 kya in the western North Pacific [83], or sudden changes in temperature during the mid-Holocene due to higher variation in temperature anomalies [84]. Fossil evidence shows 5-6 kya, when SST was ∼ 2°C higher compared to present day, reefs were prolific along the coast of Japan and found at higher latitudes [27]. This range contraction aligns well with the modeled N_e_ decline. The temperate lineage’s earlier contraction relative to the two other lineages is likely due to the temperate lineage occupying reefs (both currently and historically) at higher latitudes than the other two lineages and thus was more susceptible to cooling.

### Outlier Loci detected between *Acropora hyacinthus* lineages suggest differences in thermal tolerance

Outlier analyses revealed many potential candidate genes that could confer differential thermal tolerance between the temperate lineage relative to the two subtropical lineages. Five overlapping outlier genes were detected in both temperate-subtropical comparisons and three of these genes (two of which were annotated) have previously been implicated in the coral’s environmental stress response (ESR) [60]. These genes included centrosomal protein 290kDA and elongation of very long chain fatty acids protein 4 (EVOL4). Centrosomal protein 290kDA is involved in early and late steps in cilia formation [85] and EVOL4 is a lipid biosynthesis enzyme, which has previously been shown to exhibit differential expression between coral hosts infected with a homologous or heterologous algal symbiont strain [86]. One of the non-ESR outlier genes was helicase senataxin which is involved in DNA damage response generated by oxidative stress [87]. Variants of helicases have also previously been found in heat resistant populations of *A. digitifera* in the Ryukyus and were theorized to convey resistance against light-stress associated with heat waves [88].

Several other outlier genes were present in only one of the two temperate-subtropical lineage comparisons. Between the temperate lineage and STL lineage, ESR outlier genes included Mucin-like protein and collagen alpha-5 (VI), which have both been previously reported in the mineralizing matrices of mollusks [89]–[91]. Peroxidase was found to be an ESR outlier gene between the temperate lineage and STD lineage and several studies have linked coral peroxidase activity with anti-oxidant potential [92], [93]. Non-ESR outlier genes included Fibrillin-2, which is associated with wound healing and responds to stress in Nematostella [94] and other invertebrates [95] and was also identified as an outlier gene between *A. tenuis* lineages [96]. Hemicentin-1 – another stress response wound healing gene – was found to be an ESR outlier gene between the temperate lineage and STD lineage. This gene was also identified as an outlier between *A. tenuis* lineages [96]. Taken together, these results suggest that the temperate lineage might exhibit higher frequencies of alleles that allow for adaptation to different thermal regimes, possibly facilitating this lineage’s occupation of temperate habitats, which experience increased seasonal temperature fluctuations.

### Edge and core dynamics manifested in genomic signatures

Our genomic analyses within the temperate lineage confirm previous survey predictions [23], suggesting that the direction of expansion occurred from South to North within the temperate region – although we were unable to pinpoint the exact location of the origin. Interestingly, when exploring the assumptions of genetic consequences of range expansion we were unable to detect separation between the recently expanded Amakusa population and the range edge sites Kushimoto and Shirahama. Edge populations can share many of the same characteristics as recently expanded populations including low genetic diversity [97], increased LD [98], and increased selfing [99]. The high degree of similarity we observed might be because the Amakusa population expanded several coral generations ago (∼40 years) and continued gene flow over this time period may have dissipated the signal of a recent expansion (reviewed in [100]). This potential dissipation is corroborated by a previous study of *A. hyacinthus* population genetics in this region using microsatellites, which detected a further decrease in genetic diversity in sites that expanded more recently than Amakusa [31]. Alternatively, lack of separation between Amakusa and Kushimoto/Shirahama could point to a lasting signature of range expansion into Kushimoto/Shirahama and additional bottlenecks driving further reduction in genetic diversity in more recently expanded sites.

The observed genetic clustering between Amakusa and Shirahama/Kushimoto, despite long oceanographic distance between these regions and high migration rates from core sites predicted by larval dispersal models [31], could be driven by neutral forces or differences in thermal regimes. For the neutral force hypothesis, edge sites could be experiencing higher drift, given the evidence for asymmetrical gene flow and lower effective population sizes at these sites. However, with drift’s inherent randomness it seems unlikely that drift alone could produce such similar genetic signatures across sites on opposite sides of mainland Japan. As the three edge sites experience different thermal regimes than the two core sites, it is also possible that discrete environmental selection pressures could be driving this clustering. Indeed, mantel tests showed greater positive correlations between genetic distance and SST metrics compared to oceanographic distance, although none of these correlations were significant. This lack of significance between genetic distance and SST could be caused by high migration between Kushimoto and Shirahama, a non-linear relationship between genetic distance and temperature, or insufficient power from our current population sample size. It is possible that high migration from nearby Shirahama is driving panmixia between Kushimoto and Shirahama, however larval dispersal models predict higher migration from Kushima and Kochi [31]. Alternatively, the relationship between genetic distance and temperature is non-linear and instead there is some winter low threshold driving the separation between the core and edge sites. However, we are also limited by our small numbers of populations and effect size (i.e. Kushimoto experiences SST lower than the core sites but higher than the other two edge sites). Overall, while we cannot rule out neutral processes, evidence presented here is consistent with temperature-driven selective forces playing a structuring role in these populations, however future physiological work would need to be conducted to disentangle these hypotheses.

The presence of higher purifying selection detected at edge sites was counter-intuitive given that theoretical [6] and empirical studies [101] predict the opposite pattern. This prediction is based on the positive correlation between purifying selection and effective population size (largely due to lower inbreeding in bigger populations [102]) and the observation that deleterious allele surfing along an expansion front should lead to higher mutational load at edge populations [103]. However, there are circumstances where edge populations can demonstrate higher purifying selection. For example, bottlenecked populations can also show higher purifying selection if bottlenecks are followed by N_e_ increases [104]. Second, bottlenecks tend to purge highly deleterious mutations [105], and third, if edge and core populations occupy different environments, diverse selective constraints against *de novo* mutations could alter this signal [106]. We find the third hypothesis to be the most compelling here. Given that a large proportion of the coral genome is responsive to environmental stress, and genes that underlie CT_min_ or CT_max_ can undergo selection to remove large-effect alleles from the population by purifying selection [107], this could manifest itself as stronger purifying selection genome-wide. If a deleterious allele is more consequential in colder environments, this could explain higher purifying selection in edge populations with exposure to suboptimal temperatures leading to increased selection efficacy [108]. Future experiments elucidating thermal performance of corals in this region will help elucidate the role of purifying selection in these populations.

To begin to explore whether selective or neutral forces are important for distinguishing these genetic clusters within the temperate lineage, we performed outlier loci analyses. When investigating the outlier loci between the edge population and the core population, three ESR outlier genes were detected only one of which was annotated. This gene was protocadherin, which is a cell-adhesion protein previously implicated in pH tolerance in temperate corals relative to subtropical corals [109]. Two of the three non-ESR outlier genes were glutamate receptors, which are well-known for their importance in circadian rhythm and day/night cycles in coral [110]. These gene functions fit well into the types of environmental differences that exist across latitudes and suggest that these core and edge populations within the temperate region might be uniquely adapted to their respective environments.

## Conclusions

Our genomic analyses provide insight into the evolution and ecology of cryptic *A. hyacinthus* lineages in Japan and the mechanisms that might underly the observed population structure between edge and core populations at this species’ range edge. We found that cryptic lineages were formed in the mid-Pleistocene and have been punctuated by large increases in N_e_ that correspond with interglacial periods. Previous work has suggested that ecological opportunity from changing sea levels facilitated the adaptive radiation of Acroporids [79] and we show evidence to suggest that this also holds for intra-specific radiations as well. Within the temperate lineage we show evidence for historical occupation of marginal sites and population structure that corresponds to degrees of habitat marginalization, potentially demonstrating this lineage’s unique ability to continually expand north. Future work characterizing the physiological differences between lineages and populations will be necessary to ground truth inferences of temperature adaptation detected here. Overall, this work helps build on the growing research in this region that will be critical for evaluating the resilience of Japan’s reefs and predicting future range expansion.

## Data Availability

Raw read data can be found on the sequence read archive (SRA BioProject accession number: PRJNA735187).

## Author Contributions

SWD conceived of the project and oversaw project development. NY completed all collections including permits. JEF prepared all sequencing libraries and conducted all analyses. TY provided temperature data. JEF wrote the manuscript with contributions from SWD. All authors edited and approved the final version.

## Acknowledgements

Akira Iguchi, Takashi Nakamura, and Mikhail Matz facilitated collections, permits and shipping of samples from Japan. We acknowledge Laura Tsang, Tiffany Wong, and Brianna Regan who helped isolate all DNA samples, which was overseen by Nicola Kriefall. We are grateful to Mikhail Matz, JP Rippe, Chris Schmitt, Peter Buston, Mike Sorenson, Groves Dixon, Nicole Adams, Hanny Rivera, Sara Smith and Ekaterina Noskova for their assistance during analyses associated with the project. Thanks to Nicola Kriefall, Hannah Aichelman, Daniel Wuitchik, Colleen Bove, Lucas Fifer and Jill Grose-Fifer for comments on writing and figures. We acknowledge Boston University’s SCC who facilitated all computational work.

## Funding Statement

Funding for this project came from startup funds to SWD from Boston University and was supported by the Environment Research and Technology Development Fund (4RF 1501 and 4-1304) of the Ministry of the Environment, Japan, Grant-in-aid for young scientists (A) (17H04996). JF also received a Boston University Warren Mcleod award associated with this research.

## Declaration of interests

The authors declare no competing interests.

## Supplemental Information

**Supplemental Figure 1.**
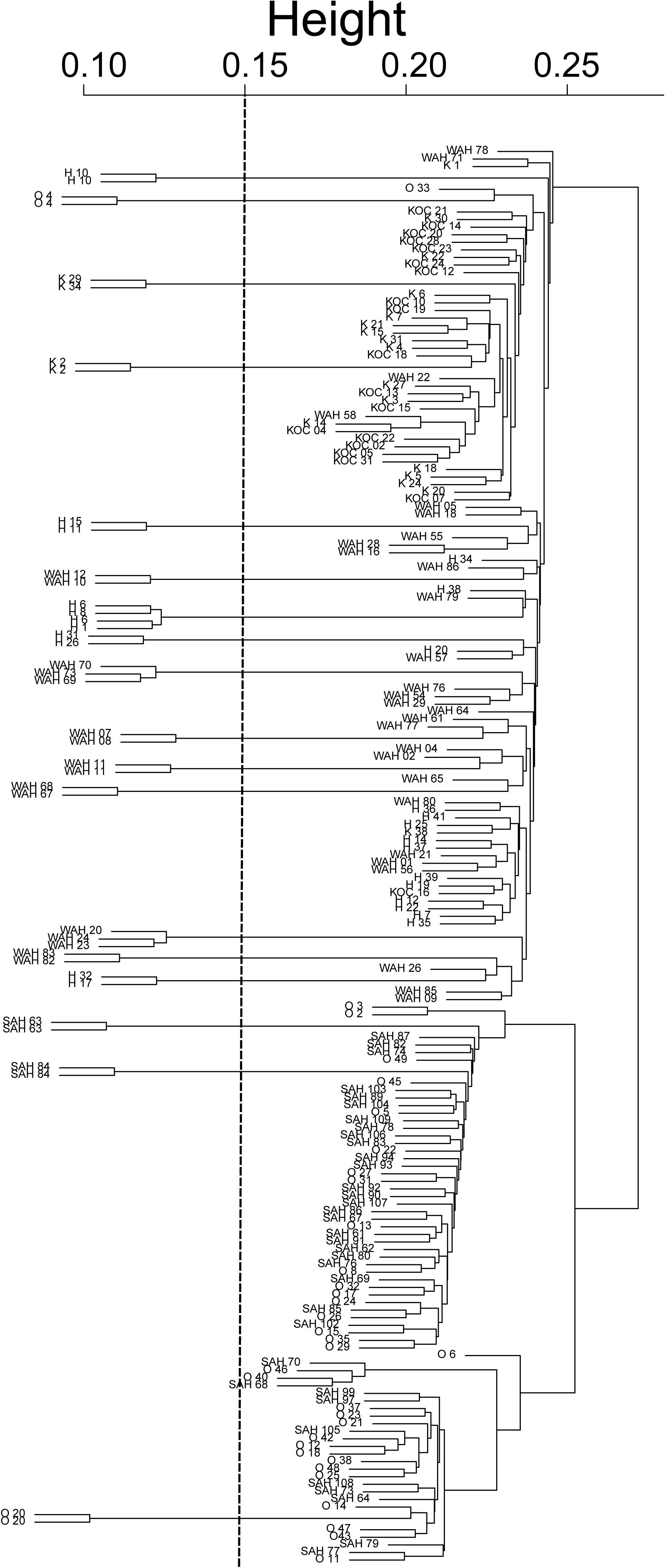
Identity by state dendrogram, dotted line represents cutoff for clone assignment.

**Supplemental Figure 2.**
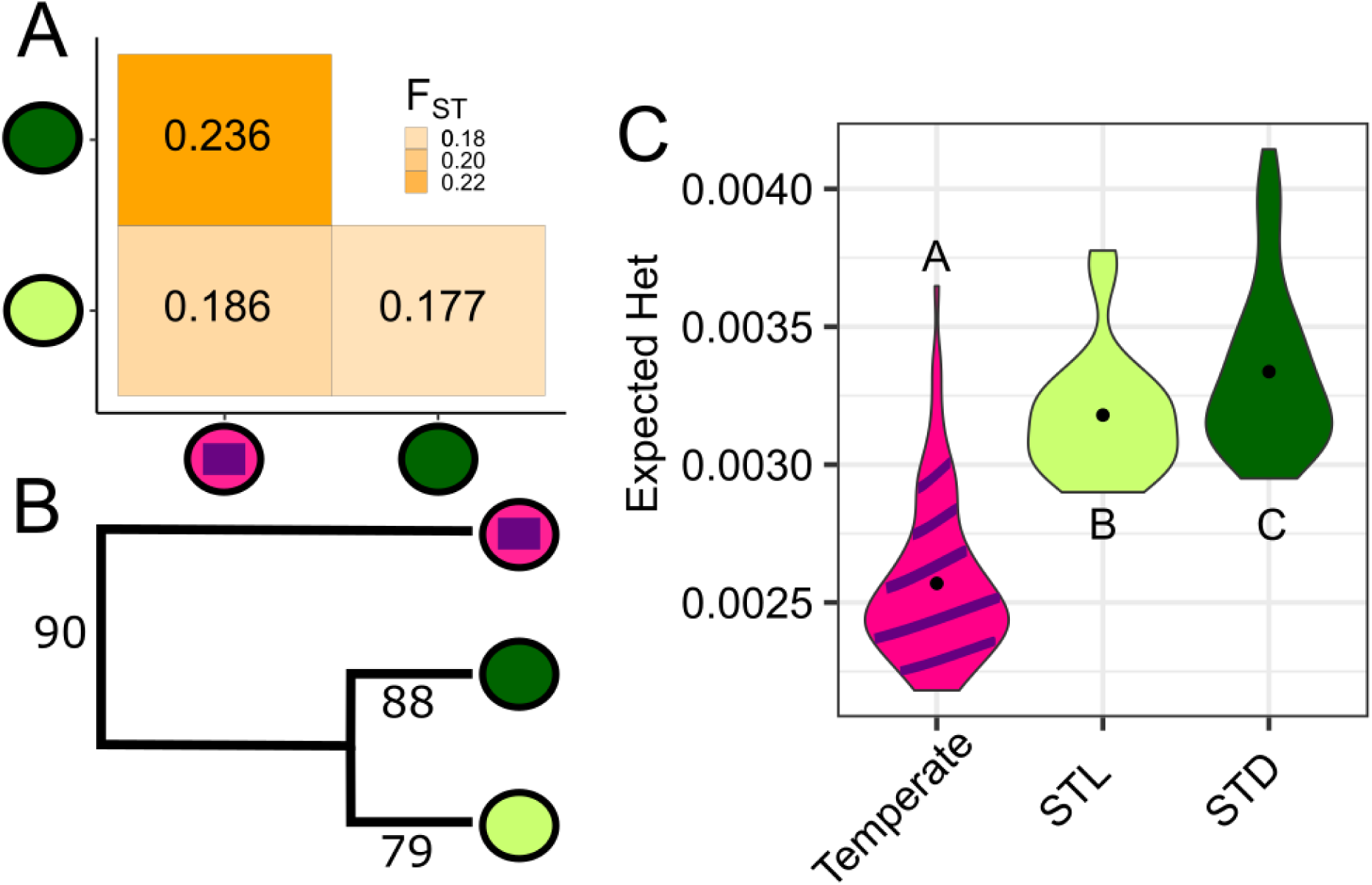
A) Pairwise global F_ST_ between all three lineages. B) RAXML tree. Numbers correspond to branch bootstrap support values. C) Expected heterozygosity estimates for each *Acropora hyacinthus* lineage. Different letters denote significant differences (p < 0.05) between lineages.

**Supplemental Figure 3.**
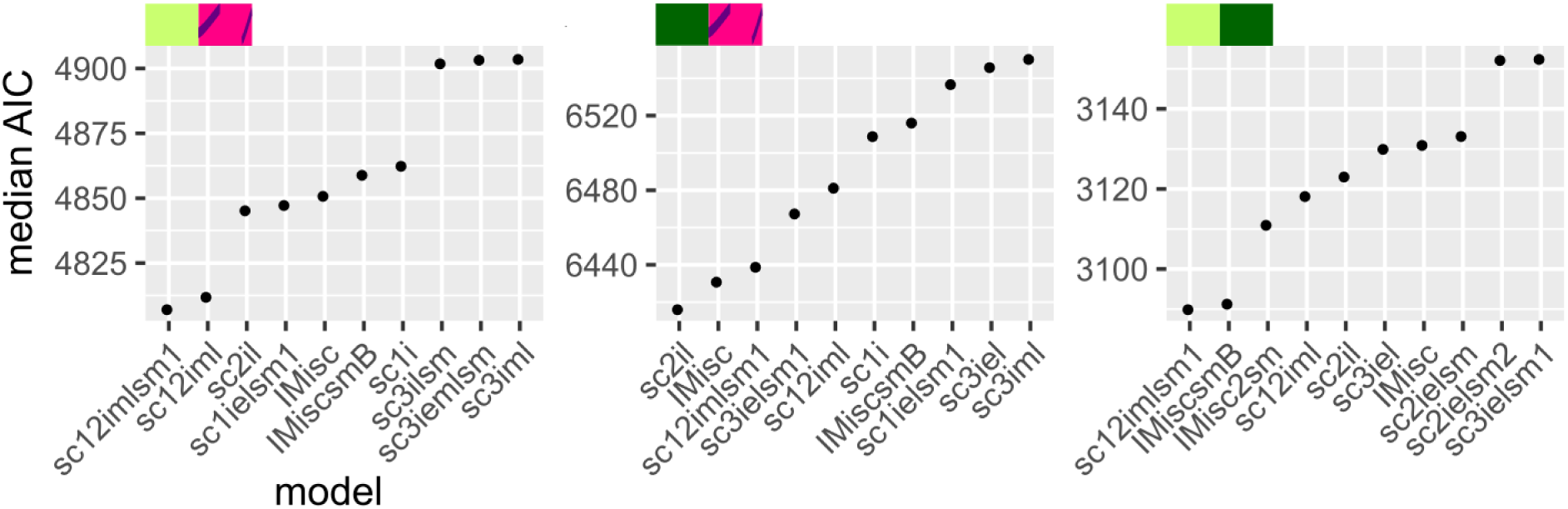
Median AIC scores for top 10 models predicted by moments. Model descriptions in Supplementary Table 2. For a complete list of all models conducted please see: https://github.com/z0on/AFS-analysis-with-moments/blob/master/multimodel_inference/moments_multimodels.xlsx

**Supplemental Figure 4.**
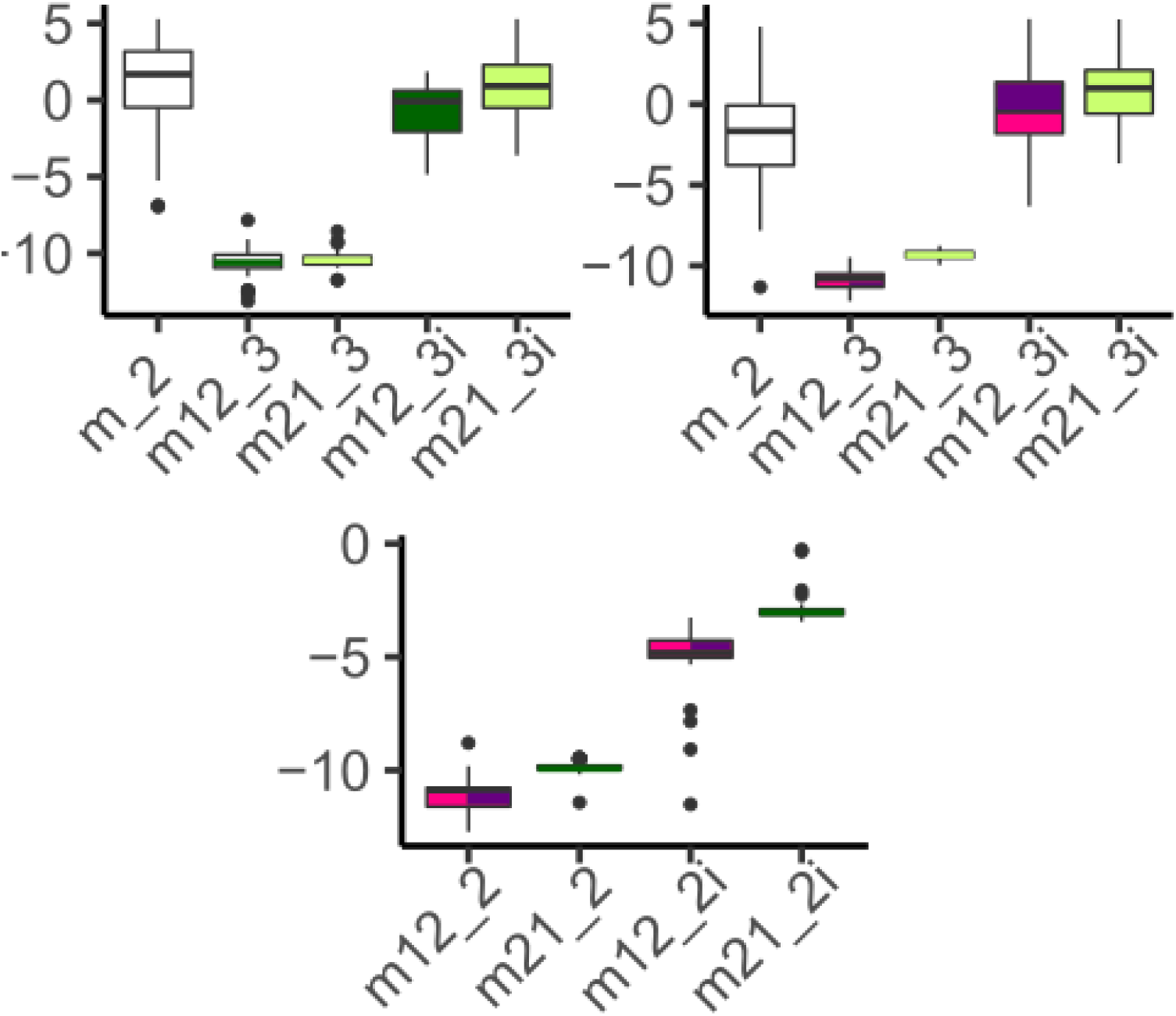
Log transformed migration rates from pairwise moments parameter output for *Acropora hyacinthus* lineage comparisons. Boxes are colored by migrant origin; i represents migration rates for genomic islands; _2 is 2nd epoch; _3 3rd epoch.

**Supplemental Figure 5.**
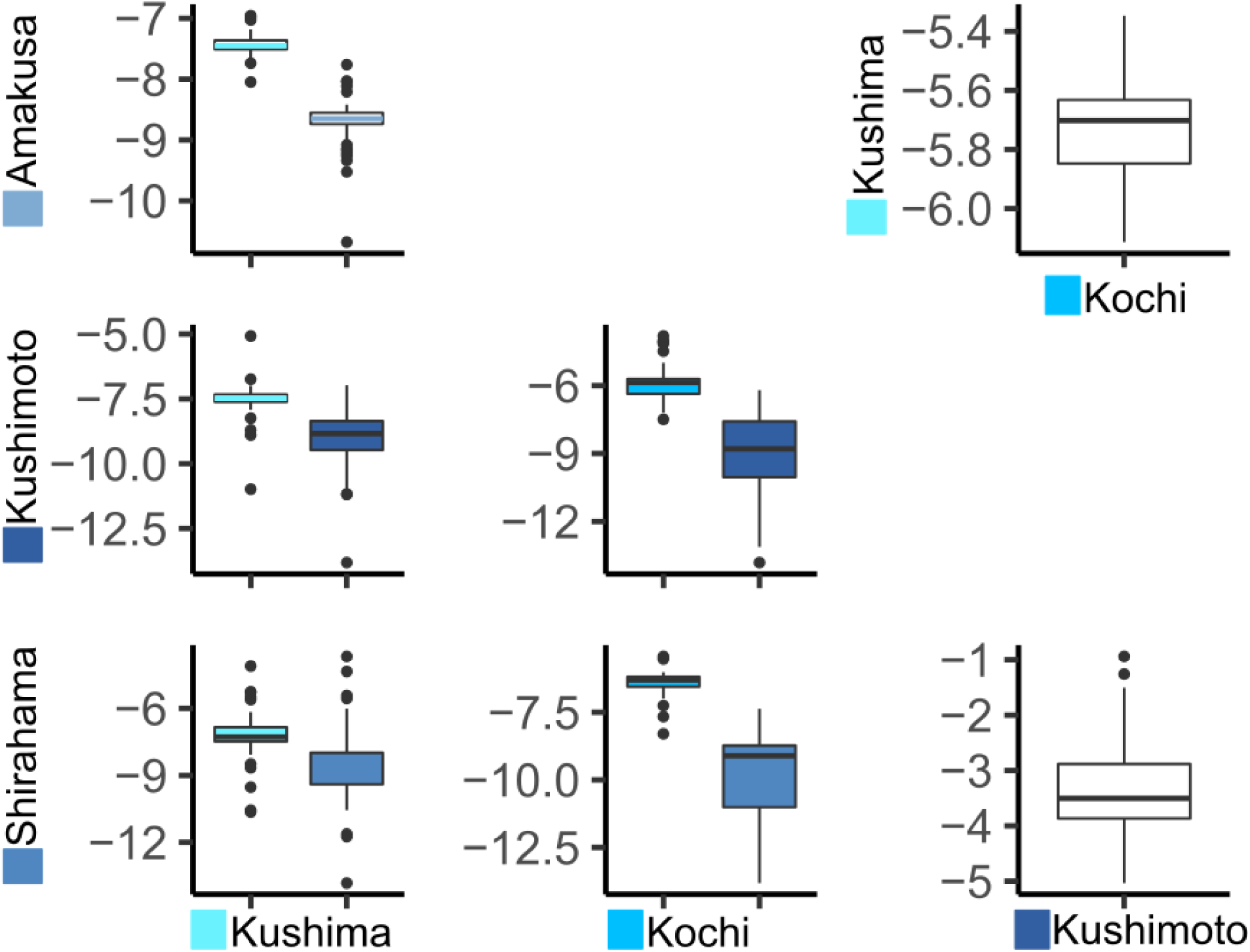
Mean migration rates for all pairwise sites within mainland Japan for the temperate *Acropora hyacinthus* lineage. Color corresponds to the source of migrants. Log transformed migration rates from pairwise moments parameter output for lineage comparisons. Boxes are colored by migrant origin; plots with single migration parameters represent symmetrical migration rates.

**Supplemental Figure 6.**
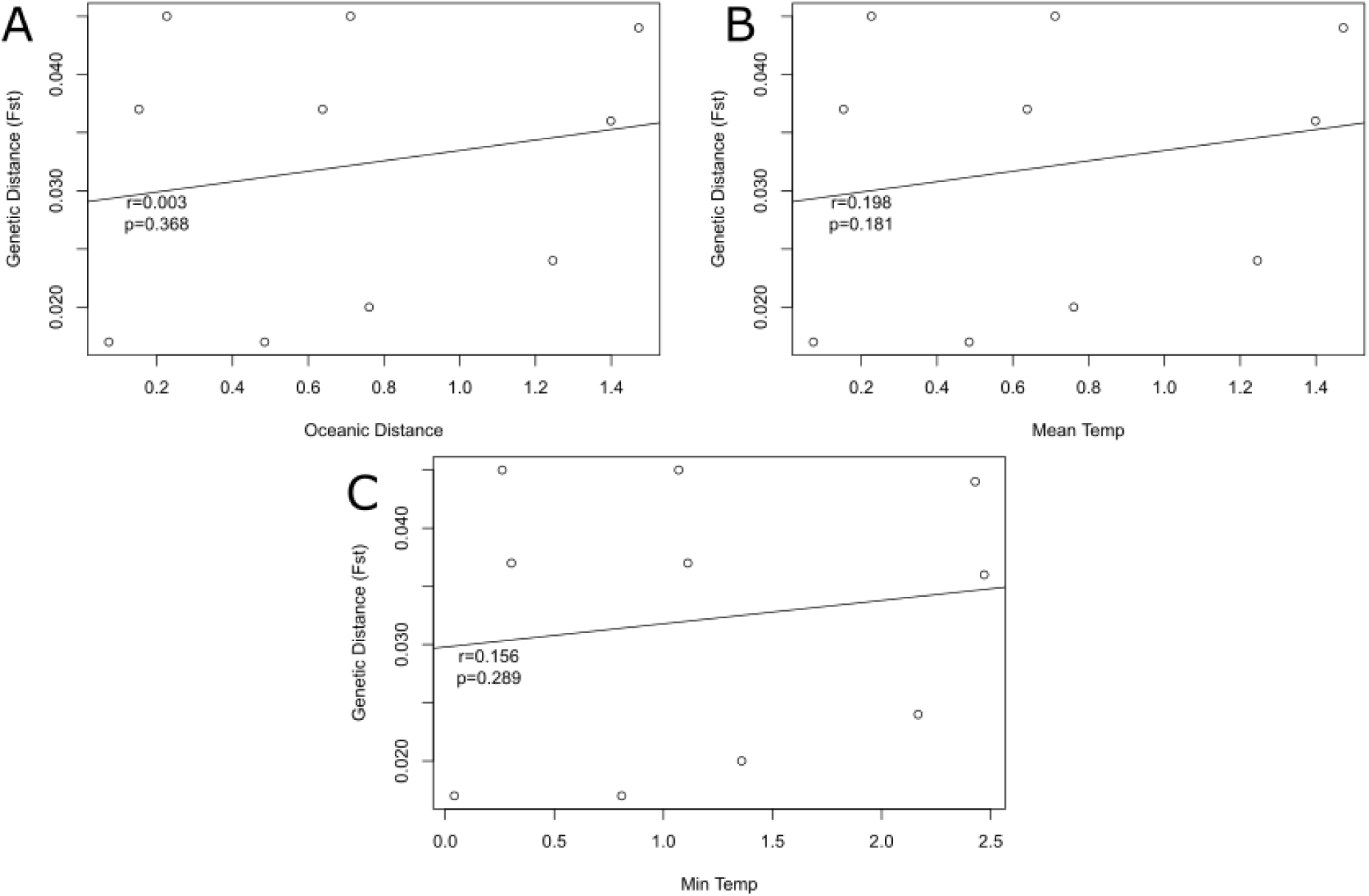
Relationships between pairwise genetic distances and A) Oceanic distances (Nakabayashi et al 2019), B) Mean monthly sea surface temperature (SST), and C) Minimum monthly SST.

**Supplemental Table 1.** Sample information for all collections.

| Site | site_lat | site_long | Depth (m) | #sampled | #sequenced | #w/o clones | Date sampled | #Temperate Lineage | #STD | #STL |
| --- | --- | --- | --- | --- | --- | --- | --- | --- | --- | --- |
| Kushima | 31.46454 | 131.2283 | NA | 38 | 21 | 20 | Dec-14 | 20 | 0 | 0 |
| Sekesei Lagoon | 24.32921 | 124.0444 | 1.0-6.6 | 51 | 40 | 37 | Jul-14 | 0 | 27 | 7 |
| Amakusa | 32.4585 | 130.1934 | NA | 42 | 26 | 18 | Sep-09 | 18 | 0 | 0 |
| Kushimoto | 33.47247 | 135.7815 | 2.4-5.5 | 31 | 23 | 17 | Jun-16 | 17 | 0 | 0 |
| Shirahama | 33.511 | 135.3482 | 0.7-9.4 | 32 | 21 | 19 | Jun-16 | 19 | 0 | 0 |
| Oura Bay | 26.53717 | 128.0579 | NA | 50 | 38 | 36 | Mar-13 | 2 | 14 | 16 |
| Kochi | 32.78871 | 132.8705 | 1.0-6.0 | 33 | 19 | 19 | Oct-11 | 19 | 0 | 0 |
| **Total** |  |  |  | **277** | **188** | **166** |  | **95** | **41** | **23** |

**Supplementary Table 2.** Model descriptions for models with top 10 median AICs in Supplementary Figure 3.

| Name | Split | Epochs | Genomic islands | Migration epochs | Symmetric migration? |
| --- | --- | --- | --- | --- | --- |
| sc12imlsm | 1 | 1 before split, then 2 | 1 | 2,3 | yes |
| sc12iml | 1 | 1 before split, then 2 | 1 | 2,3 | no |
| sc2il | 1 | 2 | 1 | 2 | no |
| sc1ielsm1 | 1 | 1 | 1 | 1 | yes in 1st epoch, no in 3d epoch |
| IMisc | 1 | 2 | 1 | 1 | no |
| IMiscsmB | 1 | 2 | 1 | 1 | yes for background genome, no for islands |
| sc1i | 1 | 1 | 1 | 1 early and late, independent | 0 |
| sc3ilsm | 1 | 3 | 1 | 3 | yes |
| sc3iemlsm | 1 | 3 | 1 | 1,2,3 | yes |
| sc3iml | 1 | 3 | 1 | 2,3 | 0 |
| sc3iel | 1 | 3 | 1 | 1,3 | 0 |
| IMisc2sm | 1 | 2, free size of 1st epoch | 1 | 1 | yes |
| sc2ielsm | 1 | 2 | 1 | 1,2 | yes |
| sc2ielsm2 | 1 | 2 | 1 | 1,2 | no in 1st epoch, yes in 2nd epoch |

Supplemental File 1

## Notes

### Competing Interest Statement

The authors have declared no competing interest.

